# Simple and Rapid High-Throughput Assay to Identify HSV-1 ICP0 Transactivation Inhibitors

**DOI:** 10.1101/2021.01.13.426532

**Authors:** Cindy Y. Ly, Chunmiao Yu, Peter McDonald, Anuradha Roy, David Johnson, David J. Davido

**Author notes:** Corresponding Author. Department of Molecular Biosciences, University of Kansas, Lawrence, Kansas, United States of America.

## Abstract

Herpes simplex virus 1 (HSV-1) is a ubiquitous virus that results in lifelong infections due to it’s ability to cycle between lytic replication and latency. As an obligate intracellular pathogen, HSV-1 exploits host cellular factors to replicate and aid in its life cycle. HSV-1 expresses infected cell protein 0 (ICP0), an immediate-early regulator, to stimulate the transcription of all classes of viral genes via its E3 ubiquitin ligase activity. Mechanisms by which ICP0 activates viral gene expression and the cellular factors involved are largely unknown. Here we report an automated, inexpensive, and rapid high-throughput approach to examine the effects of small molecule compounds on ICP0 transactivator function in cells. Two HSV-1 reporter viruses, KOS6β (wt) and *dl*x3.1-6β (ICP0-null mutant), were used to monitor ICP0 transactivation activity through the HSV-1 ICP6 promoter::*lacz* expression cassette. A ≥10-fold difference in β-galactosidase activity was observed in cells infected with KOS6β compared to *dl*x3.1-6β, demonstrating that ICP0 potently transactivates the ICP6 promoter. We established the robustness and reproducibility with a Z′- factor score of ≥0.69, an important criterium for high-throughput analyses. Approximately 19,000 structurally diverse compounds were screened and 76 potential inhibitors of the HSV-1 transactivator ICP0 were identified. We expect this assay will aid in the discovery of novel inhibitors and tools against HSV-1 ICP0. Using well-annotated compounds could identify potential novel factors and pathways that interact with ICP0 to promote HSV-1 gene expression.

## Introduction

Herpes simplex virus 1 (HSV-1) infects ~80% of the world’s population. HSV-1 is the major cause of recurrent oral-facial sores and can give rise to severe diseases such as herpes stromal keratitis and encephalitis (Roizman et al., 2007). HSV-1 cycles between two phases of its life cycle: lytic and latent infection. At first exposure, the virus productively replicates in the epithelial and fibroblast cells at the periphery and then travels along the axons of the sensory nerves that innervate these sites to establish latency in the trigeminal ganglion (Bloom, 2016). Latency is the lack of infectious virions but continued presence of the HSV-1 genome. Various stressors trigger the latent virus to lytically reactivate and may lead to recurrent symptoms, persisting as a lifelong infection. Given that HSV-1 is an obligate intracellular pathogen, cellular factors play an important role in replication and reactivation (Grinde, 2013).

Current treatments remain limited to targeting HSV-1 lytic infection and viral DNA replication. First line therapeutics include acyclic guanosine analogues such as acyclovir and valacyclovir, which upon phosphorylation by HSV thymidine kinases selectively inhibits viral DNA polymerase (Vadlapudi et al., 2013; Wilson et al., 2009). The lifelong use of these drugs has led to viral resistance (Piret and Guy, 2011). Second line therapeutics, cidofovir and foscarnet, are limited in their use due to nephrotoxicity (Wilson et al., 2009). Therefore, it is essential to identify inhibitors of novel targets that block HSV-1 lytic infection and reactivation.

We focused on infected cell protein 0 (ICP0), an immediate-early viral protein of HSV-1. ICP0 transactivates all three classes of HSV-1 genes, in part, through the destabilization and/or inhibition of host factors. ICP0 utilizes its RING-finger domain for E3 ubiquitin ligase activity targeting specific cellular proteins by conjugating them with ubiquitin, a post-translational modification (Everett, 2000; Boutell et al., 2002). Ubiquitin-mediated degradation of cellular proteins by ICP0 leads to the disruption of nuclear domain 10 (ND10), affecting cellular proliferation and differentiation, senescence, and apoptosis (Cai et al., 1993; Ching *et al.*, 2005; Zhong S *et al.*, 2000). Two ND10 constituent proteins, promyelocytic leukemia protein (PML) and Sp100, are degraded by ICP0, which inactivates the antiviral properties of ND10s (Everett *et al.*, 1998; Muller and Dejean, 1999, Lanfranca et al., 2014).

Genetic studies have shown that ICP0-null mutants are reduced for viral replication compared to wild type HSV-1 strains, demonstrating that ICP0 promotes efficient viral replication in cell culture and animal models of HSV-1 infection (Sacks and Schaffer, 1987; Leib *et al.*, 1989, Halford and Schaffer, 2000; Everett, 1989; Everett et al., 2009; Stow and Stow, 1986). Animal studies have demonstrated that ICP0 enhances the establishment of viral latency and significantly stimulates viral reactivation (Halford and Schaffer, 2001; Halford *et al*., 2006; Cai *et al*., 1993). Given this pivotal role of ICP0 in the HSV-1 life cycle, mechanisms by which ICP0 functions and the cellular pathways that it alters remain to be identified (Smith *et al.*, 2011; Hagglund and Roizman, 2004; Boutell and Everett, 2013). Genetic and cell-based assays have led to the discovery of ICP0-host interactions, but chemical biological approaches to examine these interactions have been limited.

We developed a novel approach to examine HSV-1’s ICP0 transactivator function and identify potential inhibitors of HSV-1 ICP0. Our approach utilizes two reporter viruses, KOS6β (Davido et al., 2002) and *dl*x3.1-6β (Davido et al., 2003). KOS6β (wt) and *dl*x3.1-6β (an ICP0-null mutant) have an ICP6 promoter::lacz cassette inserted between UL49 and UL50 genes. Notably, ICP0 is observed to be a potent and specific inducer of the early ICP6 gene, which encodes the large subunit of ribonucleotide reductase (Davido and Leib,1996; Davido et al., 2002; Sze and Herman, 1992; Goldstein and Weller, 1998). ICP0 transactivation activity can be monitored using a simple colorimetric-based β-galactosidase activity assay. This assay provides an inexpensive and automated high-throughput screening method. We conducted an initial screen with roscovitine, a broad inhibitor of cyclin-dependent kinases (cdks) and HSV-1 transcription, to validate the sensitivity, robustness, and reproducibility of our assay.

Our assay was used in a pilot study that screened ~19,000 compounds, and we identified 76 hits as potential ICP0 transactivator inhibitors, which included clusters of trichothecenes, lipopeptides, and cdk inhibitors. Some of the compounds in these clusters have been previously shown to impair HSV-1 replication, confirming the utility of our screen. Implications of our system are discussed.

## 2. Materials and Methods

### 2.1 Cell culture, viruses, and compounds

HepaRG cell line is derived from a liver tumor patient (Gripon *et al.*, 2002). HepaRG cells (a gift from Roger Everett) were grown in William’s E Medium containing 10% fetal bovine serum (FBS), 2mM L-glutamine, 10 U/mL penicillin, 10 U/mL Streptomycin, 50 μg/mL Insulin, and 50 μM Hydrocortisone. HepaRG cells were maintained by incubation at 37°C in 5% CO_2_. Reporter viruses, HSV-1 KOS6β (wt) and *dlx*3.1-6β (ICP0-null mutant), were used in our assays and titer as previously described (Davido et al., 2002; Davido et al., 2003). The cdk inhibitor, roscovitine, was prepared in dimethyl sulfoxide (DMSO) at a stock concentration of 50 mM (Schang *et al.*, 1998). The final concentrations of roscovitine were 50 μM and 100 μM.

### 2.2 Optimization of high-throughput assay

To optimize our assay, we examined the variables of fetal bovine serum (FBS) percentage, multiplicity of infection (MOIs), infection period, β-galactosidase assay kinetics and stability. HepaRG cells were seeded in 384-wells-plates, 25 μL of 6,750 cells per well, in phenol red-free William’s E Medium containing either 1% or 2% FBS, and incubated for 24 hours at 37°C in 5% CO_2_. Then, 10 μl of KOS-6β or *dl*x3.1-6β were added to wells at MOIs equivalent to 0, 0.2, 1, and 5. Infections proceeded for 6, 12, or 24 hours. At each time point, 10 μl of 1X lysis buffer (1% Triton X-100; 20 mM Tris-HCl [pH 8.0]; 150 mM NaCl; 1 mM dithiothreitol) was added to each well and incubated at 37°C for 20 minutes. A 4.5X β-Gal Assay buffer/CPRG solution was made with Chlorophenol Red-β-D-galactopyranoside (CPRG) (Calbiochem) and 4.5X β-Gal Assay buffer (2.475 mL 1M KCl; 19.8 mL 1M Phosphate buffer (pH 7.3); 225 uL 1M MgCl2; 1.984 mL of 14.4 M BME; H2O to 50 mL). The 4.5X β-Gal Assay buffer/CPRG solution was added to each well (10 μl/well) with a final concentration of 1 mg/mL. Absorbance was measured at 595nm at 5, 35, 80, 120, 1080 minutes post-addition of the β-Gal/CPRG solution with a PerkinElmer EnVision reader.

### 2.3 Primary screen with roscovitine

A primary screen was conducted using roscovitine (positive control) with optimized conditions (Fig. 3). HepaRG cells were seeded in 384-wells-plates and incubated for 24 hours at 37°C in 5% CO_2_. Roscovitine was then transferred into each well echo 555 acoustically (Labcyte Inc.) and preincubated for 40 minutes at 37°C in 5% CO_2_. Reporter viruses, KOS-6β and d*lx*3.1-6β were dispensed at MOI 5 and processed as described in section 2.2. For studying compound effects on cell viability, HepaRG cells were treated with roscovitine for 12 hours. An ATP-based cell viability assay was performed using Promega Cell-Titer Glo reagent according to manufacturer’s instructions. GraphPad Prism 8 was used to determine IC_50_ and CC_50_.

### 2.4 Screen with KU-HTLS libraries and cell cytotoxicity

HepaRG cells were seeded as previously described in 2.3. Each compound from the KU-HTLS was transferred by echo 555 acoustically into each well for a final concentration of 10 μM. The libraries included Selleck Bioactives, Natural Products (GreenPharma), CMLD Diversity, Analyticon Natural Products, Life Natural Products. Each compound was preincubated for 40 minutes at 37°C in 5% CO_2_. Then reporter viruses, KOS-6β and d*lx*3.1-6β, were dispensed at an MOI 5 per well, and processed as described in section 2.2. To eliminate compounds that are potential false positive hits based on cytotoxicity we used Promega Cell-Titer Glo reagent according to manufacturer’s instructions. Each compound was incubated at 10 μM for 12 hours prior to read.

### 2.5 Cycloheximide block and release

HepaRG cells were seeded as previously described in 2.3. Cycloheximide (CHX), protein synthesis inhibitor, was transferred into each well for a final concentration of 50 μg/mL and incubated for 1 hour. Then, KOS-6β and d*lx*3.1-6β were dispensed at an MOI 5 per well. After 4 hours post infection, virus and CHX was washed off with phosphate-buffer saline twice. Complete Williams E media was added to each well and the compounds were transferred by echo 555 acoustically into each well for a final concentration of 10 μM. After 20 hours of incubation, each well was processed as described in section 2.2.

### 2.6 Chemo-informatics screen

The hits from the initial screen were clustered using Canvas by Schrodinger (Schrodinger Release, 2017; Duan et al., 2010; Sastry et al., 2010). MACCS fingerprints were calculated for each compound. Compounds were clustered using hierarchical clustering and leader-follower clustering, with various merge distances/cluster radii. Leader-follower clustering of MACCS hits with a cluster radius of 0.3 yielded visually intuitive clusters.

## 3. Results

### 3.1. Assay optimization

We developed a colorimetric cell-based assay to monitor the transactivation activity of ICP0 in a 384-well plate format. The β-galactosidase viral reporter system allows for an automated colorimetric screen to rapidly process large numbers of compounds. To optimize our screen we examined several variables: serum levels, MOI, length of infectivity, and longevity of colorimetric signal, as outlined in **Figure 2**. HepaRG cells, human hepatocytes, was selected for our cell-based assay because they are easy to culture and are readily infected by HSV-1 (Everett et al., 2008). To maintain viable growth conditions for HepaRG cells and reduce possible non-specific binding of viral particles to serum, we tested 2 concentrations (1% or 2%) of fetal bovine serum (FBS) in our medium. As shown in **Figure 3**, absorbance signals for β-galactosidase activity at 1% and 2% FBS remained <0.16 in absence of virus, demonstrating no effect on β-galactosidase activity. Our assay utilizes reporter viruses, KOS6β (wt) and *dl*x3.1-6β (ICP0-null mutant), that have an ICP6 promoter::lacz expression cassette. ICP0 is a specific and potent inducer of ICP6, allowing us to utilize a β-galactosidase reporter system to monitor ICP0 transactivation activity. In presence of KOS6β or *dl*x3.1-6β, the absorbance remained consistent for either virus irrespective of MOIs or time of infectivity. As 2% FBS did not appear to impact cell viability, we decided to use media containing 2% FBS in future experiments.

**Figure 1.**
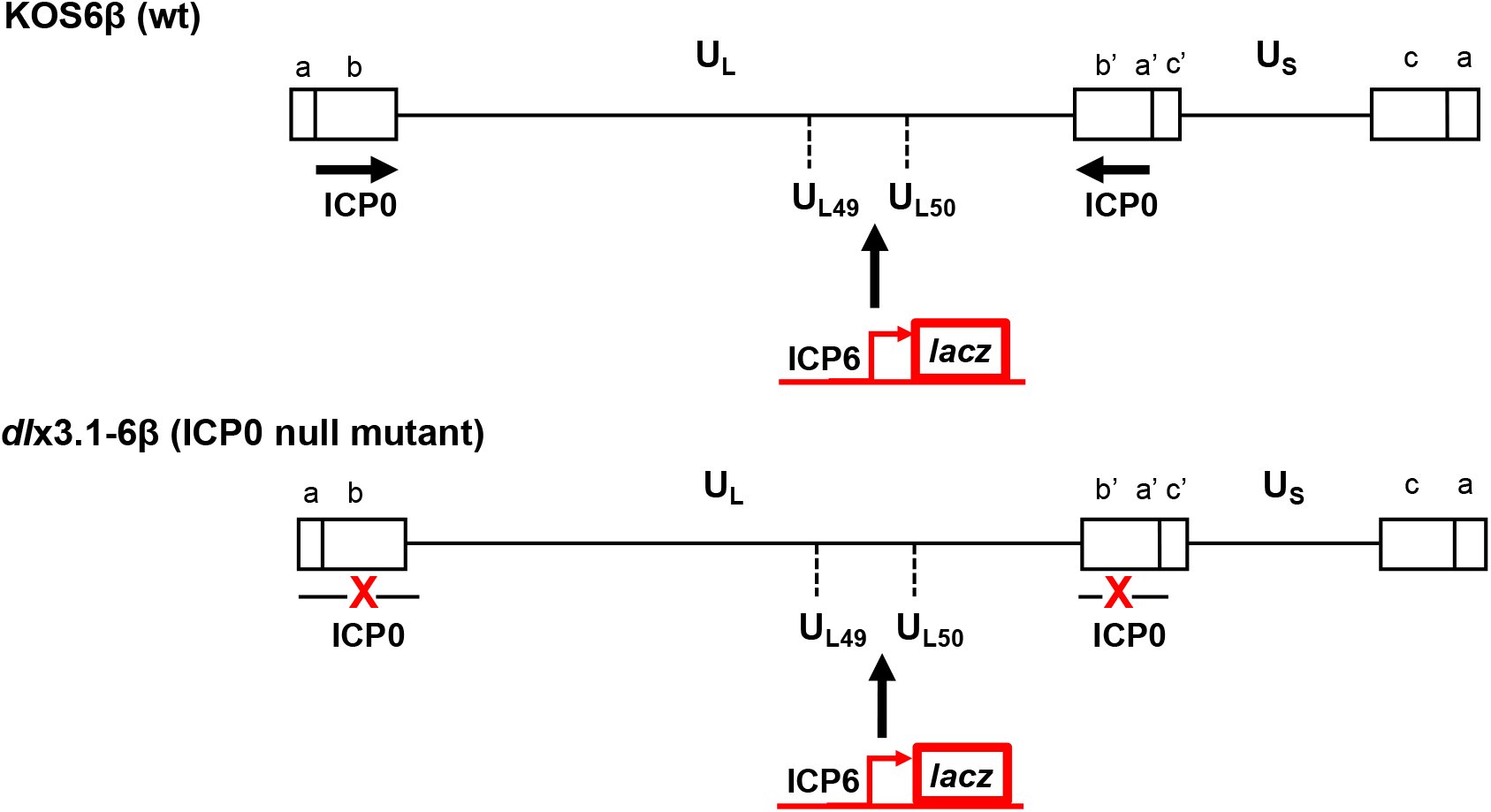
Diagram of reporter viruses. The background strain used in this approach is KOS, an HSV-1 wt strain (not drawn to scale), with unique long (UL) and unique short (US) regions of the genome flanked by an inverted repeat sequences (ab UL b’a’ and a’c’ Us ca). Two arrows represent ICP0 which contains two copies in the HSV-1 genome, and ICP0 is a specific inducer of ICP6. The reporter viruses KOS6β and *dlx*3.1-6β both contain a ICP6 promoter fused with a lacz reporter gene cassette inserted between the UL49 and UL50 genes of HSV-1. *dlx*3.1-6β, an ICP0-null mutant, has a 3.1 kb deletion in both copies of ICP0 gene. This figure is adapted from Davido, *et al*., 2003.

**Figure 2.**
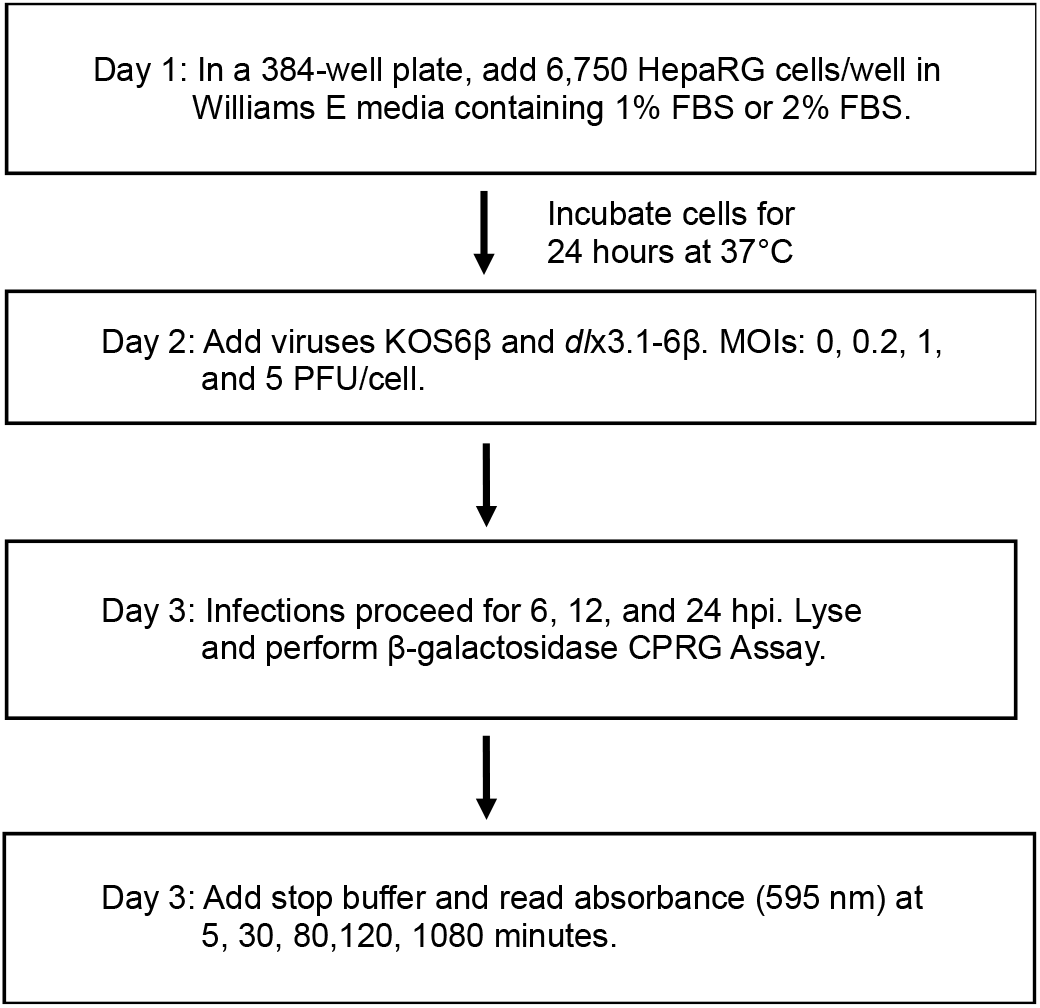
Schematic of optimization assay. Serum percentage, multiplicity of infection (MOI), infection period, and β-galactosidase stability. Schematic of optimization process: 6,750 HepaRG cells were seeded in each well of 384-well plates with 1% or 2% FBS, incubated at 37°C in 5% CO_2_ for 24 hours. Cells were then infected with KOS6β or *dl*x3.1-6β at MOIs of 0, 0.2, 1, and 5. β-galactosidase levels were analyzed at 6 hpi, 12 hpi, and 24 hpi using CPRG. β-galactosidase stability was examined at 5, 35, 80, 120, 1080 minutes. (Further description of assay is described in Methods section.)

**Figure 3.**
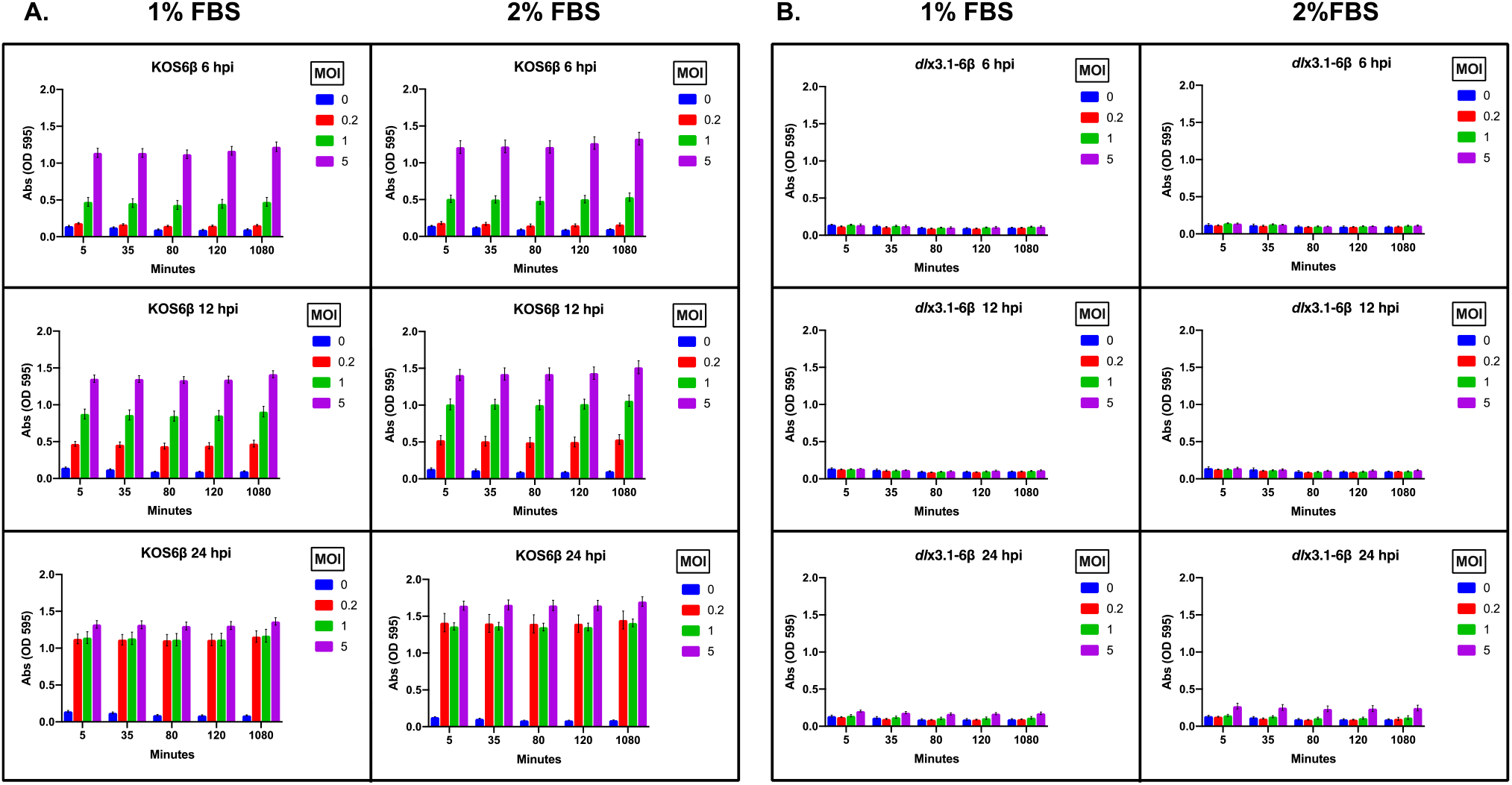
Optimization results. Histograms show optimization results of: (A) KOS6β infection and (B) *dl*x3.1-6β infection. For each reporter virus the data compares: 1% or 2% FBS, MOIs: 0, 0.2, 1, and 5, and β-galactosidase levels at 6 hpi, 12 hpi, and 24 hpi using CPRG, and assay stability from 5 to 1080 minutes.

**Figure 4.**
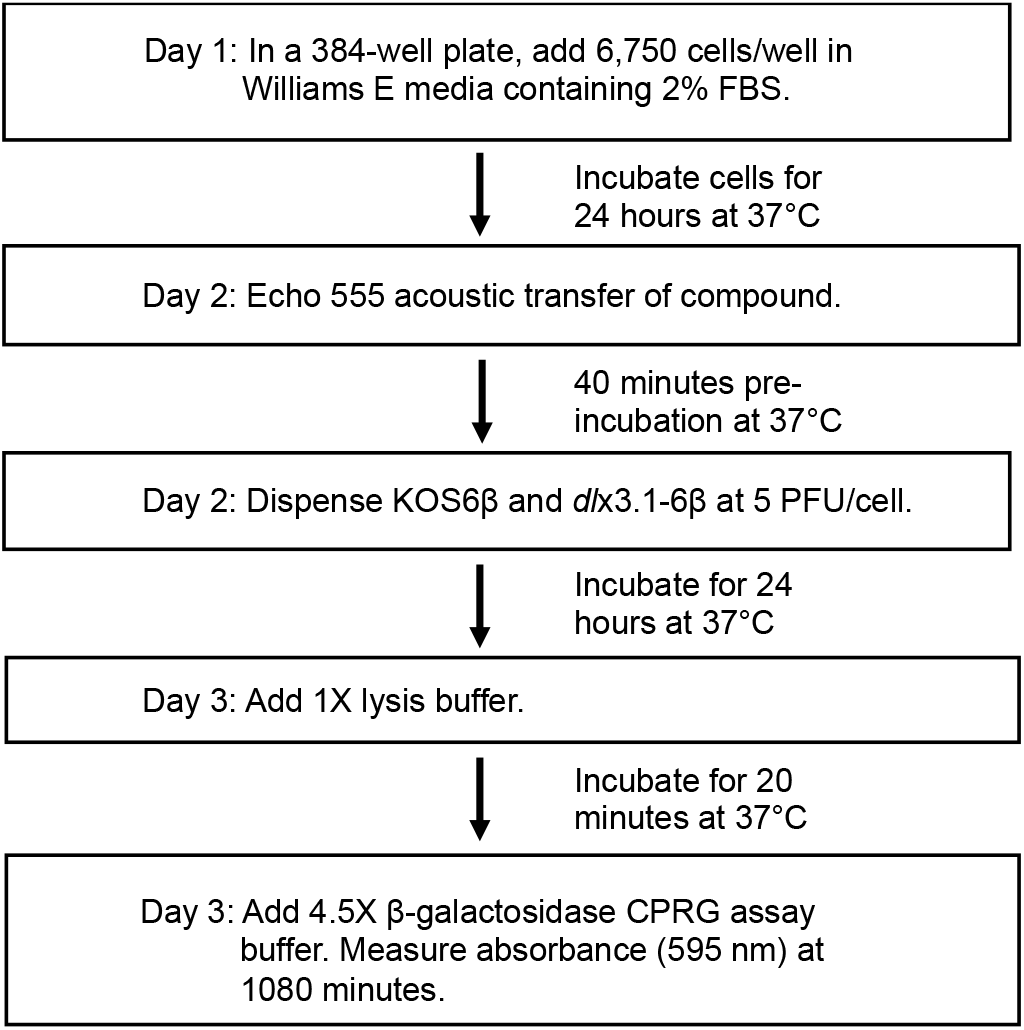
Schematic of Primary Screen. The assay used 2% of FBS, 5 PFU/cell for both reporter viruses, and 24 hpi. The approach was tested on a known small molecule inhibitor of HSV gene expression, roscovitine, as a positive control and the negative control, DMSO (0.6%). This approach was also used to screen multiple compound libraries.

We then examined MOIs at 0, 0.2, 1, and 5 PFU/cell to determine the optimal MOI for β- galactosidase activity signal. With KOS6β, absorbance signals ranged from 0.5 to 1.5, with a MOI of 5 reaching optimal signal by 24 hours post-infection (hpi). The MOIs of 0.2, 1, and 5 were clearly differentiated at 12 hours post-infection, irrespective of FBS concentration. By 24 hpi β- galactosidase levels of cells infected at MOIs of 0.2 and 1 began to approach those samples with an MOI of 5. For *dl*x3.1-6β infections at the lower MOIs (i.e., 0.2 and 1.0), the absorbance was at background levels, regardless of the infection time point. A reproducible increase in absorbance (2-3-fold) at an MOI of 5 by 24 hpi was observed compared to mock-infected cells. These data indicate that ICP0 strongly transactivates the ICP6 promoter of HSV-1, mirroring results from other published reports. We ultimately used an of MOI of 5, as it gave the highest signal relative to the background control for both reporter viruses.

The kinetics of β-galactosidase activity for all groups were analyzed at 6, 12, and 24 hpi. For KOS6β, β-galactosidase expression was noticeably detected at 6 hpi for all MOIs, with absorbance values showing marginal to substantial increases by 12 and 24 hpi. 24 hpi was selected as optimal infection time point, as maximal β-galactosidase activities were observed with the reporter viruses at an MOI of 5.

Lastly, we analyzed the stability of β-galactosidase assay after stop solution was added and absorbance read 5, 35, 80, 120, and 1080 minutes later. There appeared to be no visible differences in the absolute absorbance values between time points relative to the MOI used or time of infection. Given the stability of β-galactosidase activities, all subsequently assays were read 1080 minutes after stop solution was added. In summary, we selected the optimized conditions of 2% FBS, MOI of 5, 24 hpi, and 1080 minutes plate reads for our reporter assays.

### 3.2. Assay validation: primary screen with roscovitine

After establishing our final conditions, this assay was initially validated in a screen setting. Roscovitine, a broad cdks inhibitor, blocks the expression of many HSV-1 genes. Consequently, roscovitine was used as a positive control to validate the inhibition of HSV-1 gene expression in our reporter assay. Cells were pre-treated for 40 minutes with roscovitine over a range of concentrations. As shown in **Figure 5**, a sigmoidal-dose response was obtained for inhibition of roscovitine against the two HSV-1 reporter viruses. Roscovitine showed an inhibitory concentration of 50% (IC_50_) at 17.39 μM for KOS6β and 8.18 μM for *dl*x3.1-6β. Approximately 50% loss of cell viability was observed with 100 μM roscovitine.

**Figure 5.**
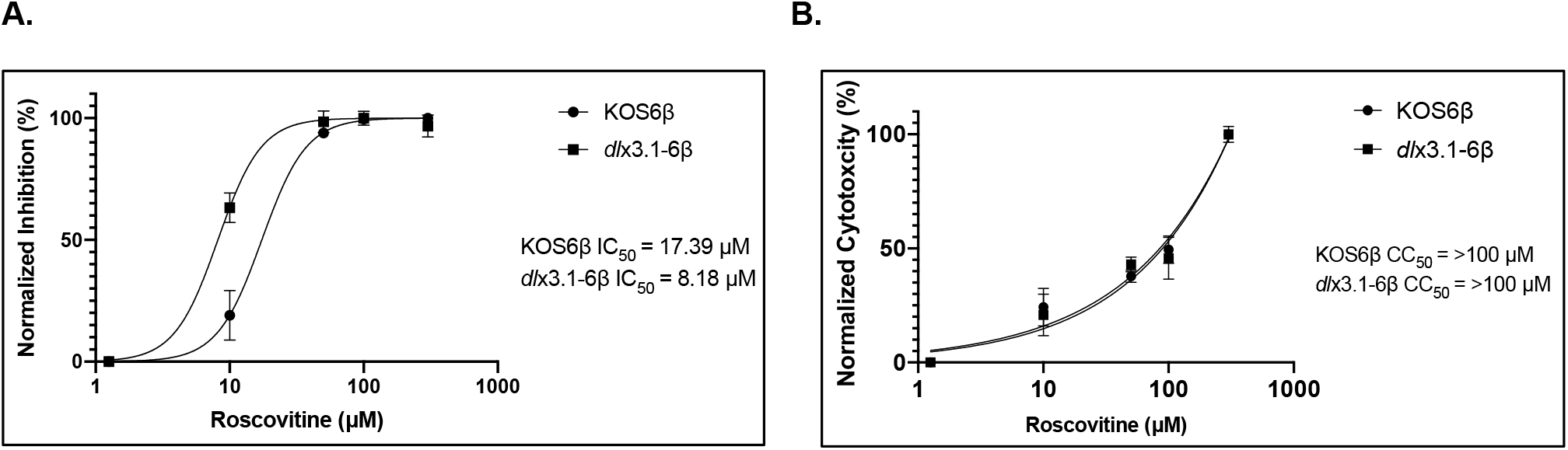
Inhibition of β-galactosidase activity from reporter viruses by roscovitine. To confirm the activity of roscovitine in our assay, dose-dependent responses to β-galactosidase expression and cytotoxicity were measured.

### 3.3. Assay robustness and reproducibility

This assay was tested for its robustness and reproducibility to be used in screening libraries of small molecule inhibitors. With KOS6β, roscovitine at 50 μM and 100 μM had a 3.75- and 9.3- fold reduction in β-galactosidase activity, respectively, compared with untreated controls (**Figure 6**). This validates the sensitivity of our assay with the use of a known small molecule inhibitor of HSV-1 gene expression. Furthermore, wells only containing the reporter virus, KOS6β, exhibited a low absorbance signal (0.16 ± 0.014), whereas wells infected with KOS6β and treated with roscovitine reached an average Abs of 0.8 ± 0.048 and 0.32 ± 0.042 at 50 μM and 100 μM concentrations, respectively. To assess the quality of the high-throughput assay the Z′, screening window coefficient was calculated (**Figure 6**). The Z′ score is a statistical indicator of assay quality, measuring assay signal dynamic range, data variation associated with sample measurement, and data variation associated with reference controls. A score between 0.5 and 1 indicates suitability of assay for high throughput screening (Zhang et al., 1999). The Z′-factor for all samples were above ≥0.69, indicative of good separation of the positive and negative controls in our assay.

**Figure 6.**
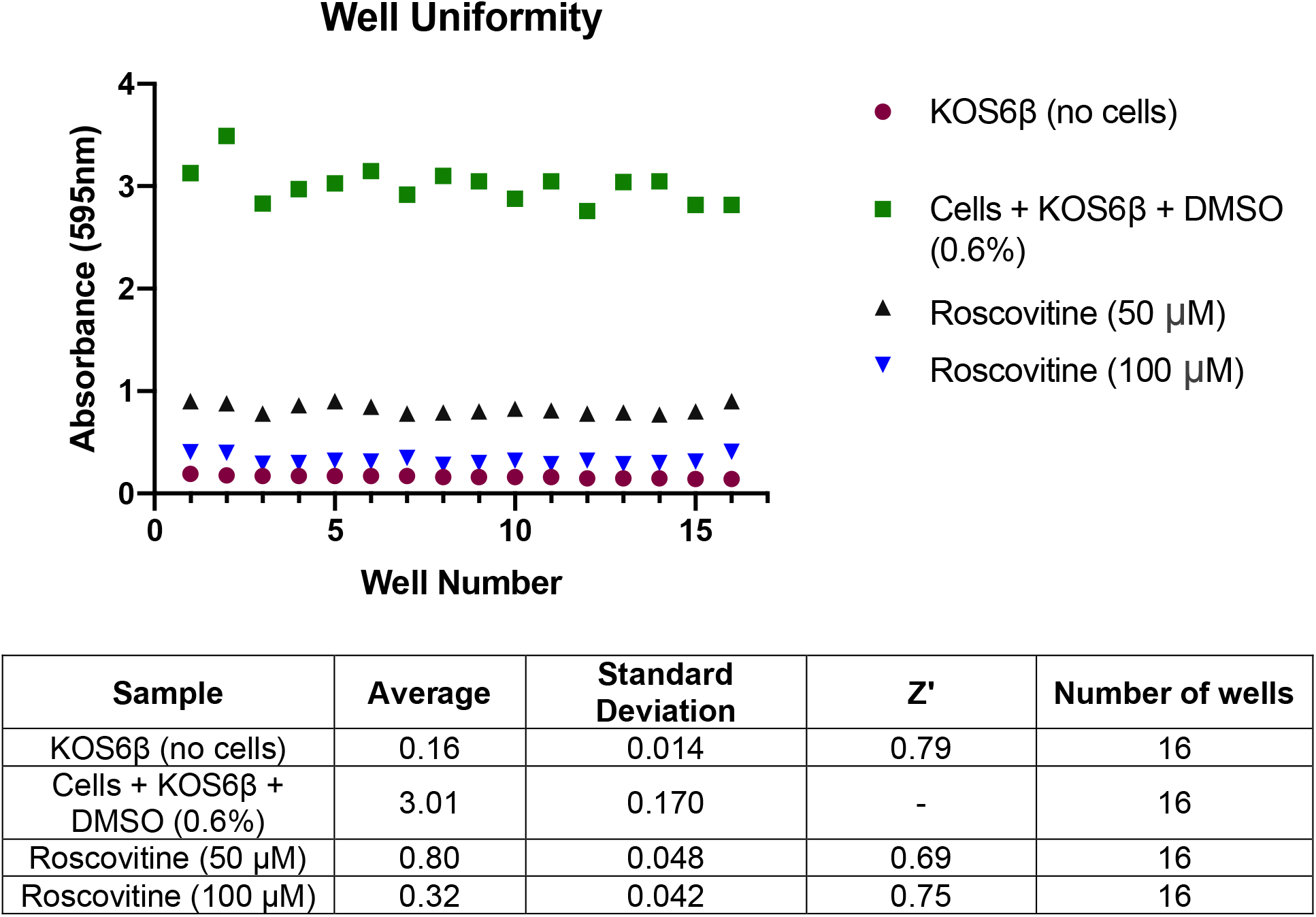
Well plate uniformity. Scatterplot of 16 wells examining in samples containing KOS6β, cells and KOS6β and DMSO (0.6%), KOS6β-infected cells plus roscovitine (50 μM), or KOS6β-infected cells plus roscovitine (100 μM). The average and standard deviation from each sample was measured to calculate the Z′ score.

### 3.4 Screen with KU-HTLS: pilot screen and secondary screen

We employed our high-throughput assay and screened ~19,000 compounds from the High Throughput Screening Laboratory at the University of Kansas (KU-HTSL). The KU-HTSL is a collection of diverse small molecules with unique scaffolds from several commercial vendors. The small molecule compounds screened included bioactive FDA approved inhibitors, natural product scaffolds amenable to chemical synthesis, purified drug-like compounds, and purified secondary metabolites. The screen of ~19,000 compounds at 10 μM resulted in a hit rate of 4.6%, 840 compounds, focusing on compounds above 3 standard deviations from the plate median. To eliminate false positives due to cytotoxicity of the compounds, ATP levels were measured for cell viability. The cytotoxicity assay reduced the number of hits to 349 compounds, hit rate of 1.9%. To help assess if the compounds are directly or indirectly blocking ICP0 transactivation activity we employed a secondary assay, a cycloheximide (CHX) block and release. CHX blocks protein synthesis and allows ICP0 transcripts to accumulate. At 4 hpi, the CHX block is released and each compound is added when ICP0 protein is expressed. The secondary assay resulted in 76 final hits, a 0.4% hit rate, which helped focus our efforts on specific compounds. We then utilized a chemo-informatic approach (Canvas by Schrodinger) to cluster compounds based on chemical structure and filter-out compounds that are potentially promiscuous or reactive. This resulted in 42 clusters, including singletons, and eliminated 6 hits flagged as pan-assay interference compounds (PAINS).

#### 3.4.1 Clusters & singletons: trichothecenes, lipopeptides, and cyclin-dependent kinases

One cluster of 10 compounds were identified to be a family of trichothecenes, secondary metabolites, produced by fungi. Trichothecenes are a class of sesquiterpenes and a few of these compounds were examined with HSV. Previous studies have shown the inhibitory effect of trichothecenes may be due to the binding of the compound to the polyribosomes, inhibiting viral protein synthesis (Tani et al., 1995; Okazaki et al., 1992; Okazaki et al., 1988). Another cluster resulted in two compounds of cyclic lipopeptides, biosurfactants produced by Bacillus subtilis. A previous study showed treatment with one of these lipopeptides reduced HSV-1 titers by >25,000-fold (Vollenbroich et al., 1997).

Several singletons, unique chemical structures, were pan-cdk inhibitors. Cellular cdks have been shown to be required for HSV-1 replication and transcription, regulating ICP0 function (Schang et al., 1998; Davido et al., 2003; Davido et al., 2002). One hit compound we identified, a known cdk-7 and −9 inhibitor, was shown to inhibit expression of all immediate-early genes including ICP0 (Hou et al., 2017). Overall, the previously studied hits (**Figure 7**) are proof of concept that our high-throughput assay is capable of identifying inhibitors of HSV-1. In future studies, we will examine the novel compounds and inhibitors identified from our library screen to determine the extent and mechanism of HSV-1 and ICP0 inhibition.

## 4. Discussion

ICP0 is a crucial viral regulatory protein that can dictate lytic infection or latency during an HSV-1 infection. We and others have demonstrated that ICP0 is a potent transactivator to all classes of HSV-1 genes, stimulating viral infection and reactivation. ICP0 is an attractive target for the development of novel antiherpetics, and such antivirals would be expected to limit HSV-1 lytic infection from reactivation. To date, identification of compounds that specifically inhibit ICP0 are limited.

To achieve this goal, we established a chemical-biological assay utilizing HSV-1 reporter viruses that allowed us to monitoring ICP0 transactivation function in tandem with small molecule compounds. Our optimization experiments led us to select the conditions of 2% FBS, MOI of 5 for the reporter viruses, 24 hpi, and measuring β-galactosidase 1080 minutes after the addition of stop solution. The assay achieved a robust and reproducible Z′-factor of ≥0.69 for various controls, which meets high throughput screening criteria. The sensitivity and effectiveness of the reporter system was validated using the broad cdk inhibitor, roscovitine, which displayed a dose-dependent response with the reporter viruses. Overall, this method was optimized to produce consistent and reproducible measurements for monitoring ICP0 transactivating activity in a high-throughput approach.

Our system described confers several advantages over other assays. One is the simplicity of the assay, where all components are added directly to the 384-well plate, requiring minimal handling of the samples and practically all steps are automated. The reporter viruses provide another advantage, as the lacz gene enables for a simple, inexpensive, and direct colorimetric screen for β-galactosidase activity. Previous assays utilized radioactive substrates or expensive equipment in fluorescent-based assays (Sekulovich *et al*., 1988); our approach eliminates those potential issues. Additionally, this assay provides a rapid and feasible method to screen multiple libraries of small molecule compounds in a 384-well plate format, which require small amounts of compounds for testing that are in micromolar range. Lastly, our high-throughput assay has the potential to be used in combination with other genetic screens (e.g., CRISPR/Cas9) to identify novel cellular factors involved in HSV-1 replication. Future work will be focused on determining how several of the 76 hits we identified in our screen inhibit HSV-1.

**Table 1.** Select hits from KU-HTLS screen. The optimized high-throughput assay was utilized to screen 19,000 diverse compounds from the KU-HTLS. After application of the primary and secondary screens, we had 76 final hits (KU-identifiers). From these hits, we identified a family of trichothecenes, two lipopeptides and several cdk inhibitors (with KU identifiers) as potent inhibitors in our assay.

| Compounds | Percent Inhibition in<br>Primary Screen | Percent Inhibition in<br>Secondary Screen |
| --- | --- | --- |
| **Trichothecenes** |  |  |
| KU0188522 | 100 | 99 |
| KU0280950 | 93 | 95 |
| KU0281790 | 100 | 99 |
| KU0281792 | 99 | 99 |
| KU0282448 | 90 | 91 |
| KU0283070 | 100 | 99 |
| KU0283235 | 98 | 99 |
| KU0283795 | 97 | 97 |
| KU0283847 | 101 | 100 |
| **Lipopeptides** |  |  |
| KU0283335 | 100 | 99 |
| KU0283824 | 100 | 94 |
| **Cyclin-Dependent Kinase Inhibitors** |  |  |
| KU0191030 | 97 | 96 |
| KU0190175 | 90 | 95 |
| KU0190205 | 94 | 96 |
| KU0189256 | 99 | 97 |
| KU0190358 | 99 | 95 |
| KU0190362 | 98 | 96 |

## 5. Acknowledgements

This work was supported by the University of Kansas (D.J.D.) and grant P20GM113117 from the National Institutes of Health (NIH) COBRE, Chemical Biology of Infectious Disease (D.J.D.). Cindy Ly was supported by the NIH Graduate Training Program in the Dynamic Aspects of Chemical Biology (T32GM008545). We thank Tom Prisinzano for assistance with chemo-informatics and members of the Davido lab for input related to this study. The content of this article is solely the responsibility of the authors and does not necessarily represent the official views of the NIH.

## References

1. Roizman B., et al. Herpes simplex viruses. Fields Virology. ed 6. Philadelphia: Lippincott Williams & Wilkins; 2013. pp. 1823–1897.

2. Bloom, DC. Alphaherpesvirus latency: a dynamic state of transcription and reactivation. Adv Virus Res. 2016;94:53–80.

3. Grinde, Bjørn. “Herpesviruses: Latency and Reactivation – Viral Strategies and Host Response.” Journal of Oral Microbiology, vol. 5, no. 1, 2013, p. 22766., doi:10.3402/jom.v5i0.22766.

4. Vadlapudi, Aswani D et al. “Update on emerging antivirals for the management of herpes simplex virus infections: a patenting perspective.” Recent patents on anti-infective drug discovery vol. 8, 1 (2013): 55–67. doi:10.2174/1574891x11308010011

5. Wilson, S et al. “Novel approaches in fighting herpes simplex virus infections.” Expert review of anti-infective therapyvol. 7, 5 (2009): 559–68. doi:10.1586/eri.09.34

6. Piret, Jocelyne, and Guy Boivin. “Resistance of herpes simplex viruses to nucleoside analogues: mechanisms, prevalence, and management.” Antimicrobial agents and chemotherapy vol. 55, 2 (2011): 459–72. doi:10.1128/AAC.00615-10

7. Smith, M., et al. “HSV-1 ICP0: Paving the Way for Viral Replication.” Future Virology, vol. 6, no. 4, 2011, pp. 421–429., doi:10.2217/fvl.11.24.

8. Everett, R D. “ICP0 induces the accumulation of colocalizing conjugated ubiquitin.” Journal of virology vol. 74, 21 (2000): 9994–10005. doi:10.1128/jvi.74.21.9994-10005.2000

9. Boutell, Chris et al. “Herpes simplex virus type 1 immediate-early protein ICP0 and is isolated RING finger domain act as ubiquitin E3 ligases in vitro.” Journal of virology vol. 76, 2 (2002): 841–50. doi:10.1128/jvi.76.2.841-850.2002

10. Cai W., et al. The herpes simplex virus type 1 regulatory protein ICP0 enhances virus replication during acute infection and reactivation from latency. J Virol. 1993;67(12):7501–7512

11. Ching RW, et al. 2005. PML bodies: a meeting place for genomic loci. J. Cell Sci. 118:847–854

12. Zhong S., et al. 2000. The transcriptional role of PML and the nuclear body. Nat. Cell Biol. 2:E85–E90

13. Zhong S., et al. 2000. Promyelocytic leukemia protein (PML) and Daxx participate in a novel nuclear pathway for apoptosis. J. Exp. Med. 191:631–640

14. Everett RD, et al. The disruption of ND10 during herpes simplex virus infection correlates with the Vmw110- and proteasome-dependent loss of several PML isoforms. J Virol. 1998;72(8):6581–6591.

15. Muller S., Dejean A. Viral immediate-early proteins abrogate the modification by SUMO-1 of PML and Sp100 proteins, correlating with nuclear body disruption. J Virol. 1999;73(6):5137–5143.

16. Lanfranca, M., et al. “HSV-1 ICP0: An E3 Ubiquitin Ligase That Counteracts Host Intrinsic and Innate Immunity.” Cells vol. 3, 2 438–54. 20 May. 2014, doi:10.3390/cells3020438

17. Sacks, W R, and P A Schaffer. “Deletion mutants in the gene encoding the herpes simplex virus type 1 immediate-early protein ICP0 exhibit impaired growth in cell culture.” Journal of virology vol. 61, 3 (1987): 829–39.

18. Leib DA, et al. Immediate-early regulatory gene mutants define different stages in the establishment and reactivation of herpes simplex virus latency. J Virol. 1989;63(2):759–768.

19. Halford WP, Schaffer PA. Optimized viral dose and transient immunosuppression enable herpes simplex virus ICP0-null mutants to establish wild-type levels of latency in vivo. J Virol. 2000;74(13):5957–5967.

20. Everett, R. D. 1989. Construction and characterization of herpes simplex virus type 1 mutants with defined lesions in immediate early gene 1. J. Gen. Virol. 70:1185–1202

21. Everett, R.D., et al. “Analysis of the functions of herpes simplex virus type 1 regulatory protein ICP0 that are critical for lytic infection and derepression of quiescent viral genomes.” Journal of virology vol. 83, 10 (2009): 4963–77. doi:10.1128/JVI.02593-08

22. Stow, N. D., and E. C. Stow. 1986. Isolation and characterization of a herpes simplex virus type 1 mutant containing a deletion within the gene encoding the immediate early polypeptide Vmw110. J. Gen. Virol. 67:2571–2585.

23. Halford WP, Schaffer PA. ICP0 is required for efficient reactivation of herpes simplex virus type 1 from neuronal latency. J Virol. 2001;75(7):3240–3249.

24. Halford WP, Weisend C, Grace J, et al. ICP0 antagonizes Stat 1-dependent repression of herpes simplex virus: implications for the regulation of viral latency. Virol J. 2006;3:44

25. Hagglund, Ryan, and Bernard Roizman. “Role of ICP0 in the strategy of conquest of the host cell by herpes simplex virus 1.” Journal of virology vol. 78, 5 (2004): 2169–78. doi:10.1128/jvi.78.5.2169-2178.2004

26. Boutell, Chris, and Roger D Everett. “Regulation of alphaherpesvirus infections by the ICP0 family of proteins.” The Journal of general virology vol. 94, Pt 3 (2013): 465–81. doi:10.1099/vir.0.048900-0

27. Davido, D. J., and D. A. Leib. 1996. Role of cis-acting sequences of the ICP0 promoter of herpes simplex virus type 1 in viral pathogenesis, latency and reactivation. J. Gen. Virol. 77:1853–1863.

27a. Davido, D. J., et al. “The Cyclin-Dependent Kinase Inhibitor Roscovitine Inhibits the Transactivating Activity and Alters the Posttranslational Modification of Herpes Simplex Virus Type 1 ICP0.” Journal of Virology, vol. 76, no. 3, 2002, pp. 1077–1088., doi:10.1128/jvi.76.3.1077-1088.2002.

28. Davido DJ, Von Zagorski WF, Maul GG, Schaffer PA. The differential requirement for cyclin-dependent kinase activities distinguishes two functions of herpes simplex virus type 1 ICP0. J Virol. 2003 Dec;77(23):12603–16. doi:10.1128/jvi.77.23.12603-12616.2003

29. Schang, L M et al. “Roscovitine, a specific inhibitor of cellular cyclin-dependent kinases, inhibits herpes simplex virus DNA synthesis in the presence of viral early proteins.” Journal of virology vol. 74, 5 (2000): 2107–20. doi:10.1128/jvi.74.5.2107-2120.2000

30. Schrödinger Release 2017-2: Canvas. 2017, Schrödinger LLC: New York, NY.

31. Duan, J., et al., Analysis and comparison of 2D fingerprints: insights into database screening performance using eight fingerprint methods. J Mol Graph Model, 2010. 29(2): p. 157–70.

32. Sastry, M., et al., Large-scale systematic analysis of 2D fingerprint methods and parameters to improve virtual screening enrichments. J Chem Inf Model, 2010. 50(5): p. 771–84.

33. Everett, Roger D et al. “Replication of ICP0-null mutant herpes simplex virus type 1 is restricted by both PML and Sp100.” Journal of virology vol. 82, 6 (2008): 2661–72. doi:10.1128/JVI.02308-07

34. Sze, P., and R. C. Herman. 1992. The herpes simplex virus type 1 ICP6 gene is regulated by a “leaky” early promoter. Virus Res. 26:141–152.

35. Goldstein, D. J., and S. K. Weller. 1988. Herpes simplex virus type 1-induced ribonucleotide reductase activity is dispensable for virus growth and DNA synthesis: isolation and characterization of an ICP6 lacZ insertion mutant. J. Virol. 62:196–205.

36. Gripon P., et al (2002). Infection of a human hepatoma cell line by hepatitis B virus. Proc. Natl. Acad. Sci. USA 99, 15655–15660.

37. Zhang, J., et al. “A Simple Statistical Parameter for Use in Evaluation and Validation of High Throughput Screening Assays.” Journal of Biomolecular Screening, vol. 4, no. 2, Apr. 1999, pp. 67–73, doi:10.1177/108705719900400206.

38. Tani N., et al. Antiviral activity of trichothecene mycotoxins (deoxynivalenol, fusarenon-X, and nivalenol) against herpes simplex virus types 1 and 2. Microbiol Immunol. 1995;39(8):635–7. doi:10.1111/j.1348-0421.1995.tb02254.x. PMID: 7494505.

39. Okazaki K., et al. Antiviral activity of metabolites of T-2 toxin against herpes simplex virus type 2. Biosci Biotechnol Biochem. 1992 Mar;56(3):523–4. doi:10.1271/bbb.56.523. PMID: 1369381.

40. Okazaki, K., et al “Growth Inhibition by Trichothecene Mycotoxins of Herpes Simplex Virus Type 2 and Its Application to Bioassay of the Toxin.” J-STAGE. 1988, pp.240–241, doi: https://doi.org/10.2520/myco1975.1988.1Supplement_240

41. Vollenbroich D., et al. Mechanism of inactivation of enveloped viruses by the biosurfactant surfactin from Bacillus subtilis. Biologicals. 1997 Sep;25(3):289–97. doi:10.1006/biol.1997.0099.

42. Schang LM, et al. Requirement for cellular cyclin-dependent kinases in herpes simplex virus replication and transcription. J Virol. 1998 Jul;72(7):5626–37. doi:10.1128/JVI.72.7.5626-5637.1998.

43. Hou, Jue et al. “Antiviral activity of PHA767491 against human herpes simplex virus in vitro and in vivo.” BMC infectious diseases vol. 17, 1 217. 20 Mar. 2017, doi:10.1186/s12879-017-2305-0

44. Sekulovich, R E et al. “The herpes simplex virus type 1 alpha protein ICP27 can act as a trans-repressor or a trans-activator in combination with ICP4 and ICP0.” Journal of virology vol. 62, 12 (1988): 4510–22.

